# Deletion of 12-lipoxygenase normalizes platelet function after storage and transfusion in thrombocytopenic mice

**DOI:** 10.1101/2022.09.23.509265

**Authors:** Hannah J. Larsen, Daire Byrne, S. Lawrence Bailey, Massiel C. Stolla, Michael Holinstat, Xiaoyun Fu, Moritz Stolla

## Abstract

**Objective:** Platelets for transfusion are stored for 5-7 days. During storage, platelets undergo numerous detrimental functional changes. In the current study, we sought to understand how genetic deletion of 12 –lipoxygenase (12-LOX) affects platelets during storage, before, and after transfusion.

**Approach and Results:** We obtained platelets from wild-type (WT) and 12-LOX-/-mice and performed storage studies for 24 and 48 hours. Using LC-MS/MS-MRM, we showed that ω-3 and ω-6 fatty acids increased significantly in stored platelets from 12-LOX-/-mice, while oxylipins were significantly lower than in WT platelets. The circulation time of fresh 12-LOX-/-platelets was significantly shorter than that of fresh WT platelets, but no differences were observed after storage. Baseline αIIbβ_3_ integrin activation was significantly lower before and after 24 hours of storage in 12-LOX-/-platelets than in WT platelets. Surprisingly, after transfusion, we observed more baseline αIIbβ3 integrin activation in 12-LOX-/-platelets than in WT platelets. In line with this, transfusion of stored 12-LOX-/-platelets led to more frequent and significantly faster vessel occlusions than transfusion of stored WT platelets in a FeCl_3_-induced carotid artery injury model in thrombocytopenic mice.

**Conclusion:** Deleting 12-LOX improves the post-transfusion function of stored murine platelets. Pharmacologic inhibition of 12-LOX or dietary alterations of ω-3 and ω-6 PUFAs could significantly enhance human platelet quality and function after storage. Future studies must determine the feasibility and safety of 12-LOX inhibition in stored and transfused human platelets.

## Introduction

Lipoxygenases (LOXs) are a family of dioxygenases that catalyze the subtraction of hydrogen atoms from polyunsaturated fatty acids (PUFAs) followed by addition of dioxygen.^1^ LOXs constitute key pro-oxidative enzymes in cellular redox homeostasis.^2, 3^ During platelet activation, cytosolic phospholipase A2α (cPLA2α) hydrolyzes phospholipids and releases arachidonic acid (AA) from the plasma membrane.^4^ The platelet-restricted 12-lipoxygenase (12-LOX) oxygenates AA, to form the oxylipin 12-hydroperoxyeicosatetraenoic acid (12-HpETE), which is quickly metabolized to 12-hydroxyeicosatetraenoic acid (12-HETE).^5^ 12-HETE is released from platelets and can activate and recruit platelets in a similar fashion as other second wave mediators.^6, 7^ Notably, 12-HETE and other HETE levels in stored platelets have negatively correlated with platelet survival upon transfusion in humans.^8^ Storage of platelets is necessary to make them available for thrombocytopenic or bleeding patients. For this purpose, platelets are stored at room temperature for up to 5-7 days. During storage, platelets undergo numerous detrimental changes collectively called the “storage lesion,” which causes reduced platelet survival and function. The fact that HETEs negatively correlate with platelet performance *in vivo* suggests that the storage lesion detriment is partially due to bioactive lipid mediators, including oxylipins.

The storage lesion has recently gained attention due to the surprising outcomes of clinical platelet transfusion trials. In two trials, stored platelets worsened outcomes and increased bleeding in neonates and patients with intracranial hemorrhage on antiplatelet therapy.^9, 10^ Accumulation of inflammatory lipid mediators and loss of platelet function during platelet storage are possible explanations for these findings. HETEs accumulate during platelet storage and lead to functional sequelae. Infusion of HETEs caused transfusion-associated acute lung injury (TRALI) in rats.^11^ Similarly, injection of 12-HETE into the skin of humans causes neutrophil infiltration.^12^ In contrast, no neutrophil accumulation was observed in the lungs when stored platelets with elevated 12-HETE levels were transfused to healthy humans, hinting at the necessity of additional priming events to cause lung inflammation.^13^

While previous studies highlight the importance of HETEs, they are not the only oxylipins of importance for platelets. Besides AA, 12-LOX (and other LOXs) also metabolizes other PUFAs into oxylipins which regulate and alter platelet function.^14–17^ Mice injected with 12-hydroxyeicosatrienoic acid (12-HETrE), an oxylipin derived from dihomo-gamma-linolenic acid (DGLA), showed reduced integrin activation and reduced α-granule secretion.^18^ Similarly, in mice injected with 14-hydroxydocosahexaenoic acid (14-HDoHE) or 11-HDoHE, oxylipins derived from docosahexaenoic acid (DHA), showed similar results.^19^ In addition, oxylipins are important mediators of endothelial function.^18^

In the current study, we investigated how genetic deletion of 12-LOX affects the accumulation of metabolites in the storage bag and stored platelet function *in vitro* and *in vivo*. We hypothesized that deletion of 12-LOX prolongs platelet *in vivo* survival and reduces platelet function.

## Methods

### Reagents

ACD-A was purchased from Fenwal Inc., Lake Zurich, Illinois. Leo.F2 anti GPIIbIIIa, CD41/CD61 (Dylight 649, M025-3),Wug.E9 anti CD62P (FITC, M130-1), and JON/A anti high affinity conformation GPIIbIIIa, CD41/CD61 (PE, M023-2) were purchased from emfret Analytics, Wurzburg Germany. Convulxin (ALX-350-100-C050) was purchased from Enzo Life Sciences, Inc., Farmingdale, New York. PAR-4 Agonist peptide (RP11529) was purchased from GenScript, Piscataway, New Jersey. Arachidonic Acid (P/N 390) and ADP (P/N 384) were purchased from Chrono-Log Corporation, Havertown, Pennsylvania. NHS-biotin, Ferric Chloride (FeCl_3_) (anhydrous, reagent grade) was purchased from Carolina Biological Supply Company, Burlington, North Carolina. Saline (0.9% sodium chloride, injection, USP) was purchased from ICU Medical, Inc., Lake Forest, Illinois. hIL4R-TG depletion antibody (Human IL-4R alpha Antibody, MAB230) was purchased from BioTechne R&D Systems, Minneapolis, Minnesota. Heparinized capillary tubes (41B2501) were purchased from Kimble, Rockwood, Tennessee. Modified tyrode’s buffer was prepared as previously described (137mM NaCl, 0.3mM Na2HPO4, 2mM KCl, 12mM NaHCO3, 5mM HEPES, 5mM glucose, pH 7.3) containing 0.35% BSA and 1mM CaCl2.^20^

PUFAs, Oxylipins, and derterium-labeled internal standards were purchased from Cayman chemical. J.T. Baker LC/MS-grade acetonitrileand methanol were purchased from VWR. Optima LC/MS-grade 2-propanol was purchased from Fisher Scientific. LC/MS-grade ammonium acetate was purchased from Sigma-Aldrich.

### Mice

12-LOX-/-mice were provided by Michael Holinstat. C57BL/6J (wild type, strain number 000664) mice were purchased from the Jackson Laboratory and were bred in house. hIL4R-TG mice were provided by Jerry Ware and bred in house.

### Sample preparation and platelet storage with function testing pre and post-transfusion

We collected whole blood from WT or 12-LOX-/-mice in acid citrate dextrose solution A (ACD-A) via retroorbital bleeding using heparinized capillary tubes. Platelet-rich plasma (PRP) was isolated from whole blood and stored at RT (22 °C) with continuous agitation for 24 hours. For pre-transfusion testing, we sampled platelets fresh and after storage. We washed the platelets by adding 0.5uM PGI_2_, spinning at 3200 rpm for 5 minutes, and resuspending in murine Tyrode’s Buffer. Washed platelets were adjusted to a platelet count of 300×10^3^/μL, and platelet function was tested by flow cytometry. Platelets were identified by forward, and side scatter as Leo.F2 positive. The samples were stimulated with different agonists (PAR-4 Peptide 0.5mM, Convulxin 100ng/mL, ADP 5uM, and Arachidonic Acid 0.5mM) and analyzed for mouse aIIbb3 activation using Jon/A PE antibody and measured for α-degranulation by an anti-CD62P FITC antibody.

For transfusion experiments, we adjusted the platelet concentration to 200 ×10^3^/μL and transfused 500uL into platelet-depleted hIL4R-TG mice (see below). After the transfusion, heparinized blood samples were collected from each recipient at 1, 4, and 24 hours. The transfused platelet population was identified by forward, side scatter, and as Leo.F2 positive. The samples were stimulated with different agonists (ADP 5uM, Convulxin 100ng/mL, and PAR-4 Peptide 0.5mM) and analyzed for activation of mouse aIIbb3 using a Jon/A-PE antibody to measure αIIbβ3 integrin activation level by flow cytometry.

### hIL4R-TG Thrombocytopenia Assay

hIL4R-TG animals were treated with an anti-hIL4 antibody (1.5-2.5 μg/g body weight) between 12 and 24 hours before transfusion. We confirmed thrombocytopenia before transfusion by verifying that the platelet count was less than 10% of the baseline platelet count pre-antibody treatment and tested for platelet counts 1 and 24 hours after antibody treatment. For *in vivo* surgery studies, we verified that the platelet count was less than 10% of the baseline platelet count after 12 hours.

### Liquid Chromatography-Tandem Mass Spectrometry (LC-MS/MS)

As described previously, free PUFAs and oxylipins in platelets were analyzed by liquid chromatography-tandem mass spectrometry (LC-MS/MS).^21–23^ Briefly, analytes were extracted by 80% methanol (vol/vol) with an internal standard mixture. Liquid chromatography-tandem mass spectrometry (LC-MS/MS) analysis was performed using a mass spectrometer (QTrap 6500, AB Sciex) coupled with an ultra-performance liquid chromatographer [UPLC] (Acquity I-Class, Waters). Analytes were separated on a C18 column (Acquity HSS T3, 2.1 × 100 mm, 1.8 μm, Waters). The mobile phase was composed of (A) water/acetonitrile (95/5, vol/vol) with 5 mmol/L ammonium acetate and (B) 2-propanol/acetonitrile/water (50/45/5, vol/vol/vol) with 5 mmol/L ammonium acetate. Analytes were detected using multiple reaction monitoring (MRM) in the negative ion mode and were quantified by peak area relative to their deuterium-labeled analogs. Data were collected and processed using computer software (Analyst Version 1.6.2, MultiQuant Version 2.1.1, AB Sciex).

### *In vivo* biotin platelet labeling

To test trace 12-LOX-/- and WT platelets, we used in vivo biotinylation of platelets. We used wild-type (WT) mouse platelets and biotinylated with N-hydroxysuccimido (NHS) biotin based on a previously published protocol.^24^ In brief, we injected WT and 12-LOX-/-mice with 1-2 mg of NHS biotin and obtained whole blood after approximately 4 hours. We generated PRP and injected WT mice with 200μL biotinylated platelets from either 12-LOX-/- or WT mice and platelets were traced over multiple time points in a 3-4 day window by flow cytometry and inputted into a regression model. To validate this method, we injected GFP+ platelets from UbIC-GFP mice with *in vivo* biotinylated platelets and compared their survival in WT mice, observing no significant differences between the two groups.

### *In vivo* platelet function assay

Platelets were stored as described above and washed before transfusion into platelet-depleted hIL4R-TG mice (see description above). Mice were anesthesized with isoflurane (5% for induction and 1.5% for maintenance in 1.0 L/min oxygen).^25^ FeCl_3_ injury of the carotid artery was performed as described by us and others before.^26^ In brief, the left carotid artery of an adult male mouse (8-10 weeks of age) was exposed to a 1mm × 2mm strip of Whatman qualitative filter paper, saturated with FeCl_3_ (2M in PBS) for 4 minutes, then rinsed with PBS. The arterial flow rate was recorded for 30 minutes with a Doppler flow probe (Transonic, Ithaca, NY). The time to occlusion was measured as time from filter paper application to complete cessation of blood flow (when flow was equal to 0.0 mL/min), which had to remain stable for the 30 minute duration of the assay. The area under the curve was calculated (Labchart Pro 8, AD Instruments, Dunedin, Otago, New Zealand). The mice were sacrificed at the end of the procedure.

### Statistical analysis

Results are reported as mean ± SEM, and statistical significance was assessed by unpaired 2-tailed Student t-test, with Holm-Sidak adjustment for multiple comparisons as necessary. A p-value less or equal to 0.05 was considered significant.

## Results

### Metabolomics of wild-type and 12-LOX-/-platelets during storage

We tested platelets from WT and 12-LOX-/-mice at baseline, after 24, and after 48 hours of storage by targeted metabolomics using LC-MS/MS-MRM. We included ten ω-6 and ω-3 polyunsaturated fatty acids (PUFAs) and 28 oxylipin markers from the lipid metabolism (Figure 1A). Overall, PUFA levels did not differ between fresh WT and 12-LOX-/-platelets, except for eicosapentaenoic acid (EPA) and docosapentaenoic acid (DPA), which were higher in fresh 12-LOX-/-platelets. However, after storage, almost all tested PUFAs increased significantly in 12-LOX-/-platelets compared to WT platelets (Figure 1A, C). In contrast, the vast majority of oxylipins were significantly lower in 12-LOX-/-platelets compared to WT, including the critical lipid mediator 12-HETE (Figure 1D). Because 12-LOX can use several PUFAs as substrates, other oxylipins were significantly decreased in 12-LOX-/-platelets, including 14-HDoHE from DHA and 12-HEPE from EPA. Additionally, there was a significant decrease in 15-HETE, 17-HDoHE, and 13-HODE in the 12-LOX-/-platelets compared to WT. We also observed a significant decrease in metabolites mediated by the cyclooxygenase (COX) pathway, including PGE2, PGD2, thromboxane B2 (Supplemental Figure 1B), and 12-HHTrE. Arachidonic acid, a known platelet agonist, increased significantly in both WT and 12-LOX-/-platelets, but the levels were significantly higher in 12-LOX-/-platelets at both 24 and 48 hours of storage (Figure 1E). To test whether AA accumulated in the storage supernatant, we analyzed platelet-poor plasma from stored platelets. Indeed, we found AA levels to be significantly higher in the supernatant (Supplemental Figure 1A). The endproduct of the AA metabolism through COX-1 in platelets is the second-wave mediator thromboxane A2. In contrast, we found a trend for lower thromboxane B2 (the stable product of A2) levels in the supernatant of 12-LOX-/-platelets (Supplemental Figure 1B). The ω-6 to ω-3 fatty acid ratio was significantly lower in 12-LOX-/-than in WT platelets at baseline but further decreased markedly after 24 hours and remained stable after 48 hours of storage (Figure 1F). After storage, the absolute ω-3 and ω-6 PUFAs increased significantly in 12-LOX-/-platelets compared to WT platelets. Therefore, the decreased ω-6/ ω-3 ratio indicates a pronounced increase in ω-3 PUFAs compared to ω-6 PUFAs. In contrast, the ratio remained stable in stored WT platelets (Figure 1F).

**Figure 1:**
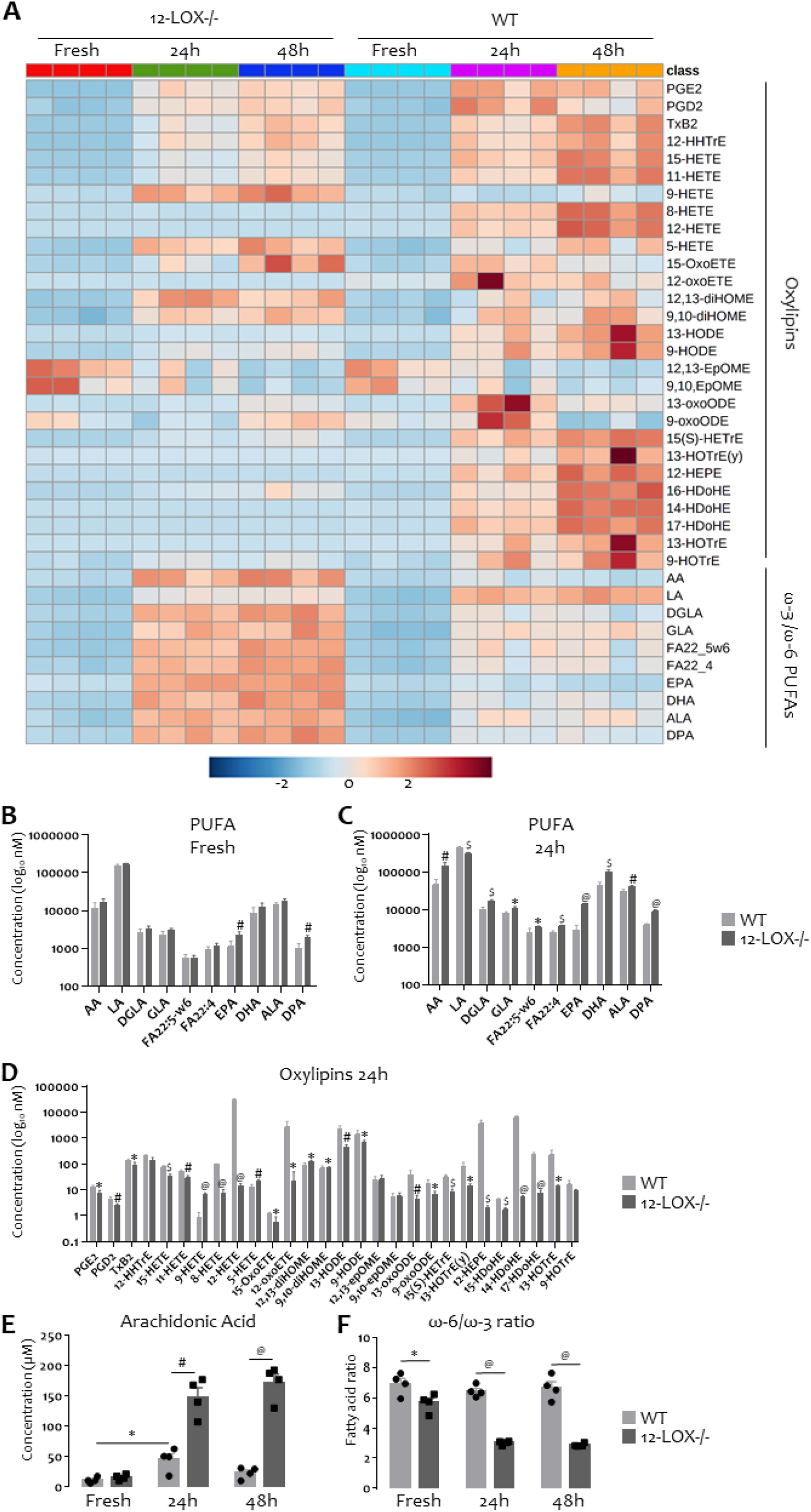
Targeted lipidomics of 12-LOX-/- and WT platelets.

Platelets were isolated and tested fresh or after 24 hours and 48 hours of storage. **(A)** Heatmap of 12-LOX-/- (left), tested fresh (red), 24h storage (green), and after 48h storage (blue). WT (right), fresh (light blue), 24h stored (magenta), and 48h stored (orange). **(B)** Individual PUFA concentrations tested fresh in WT (light grey bars) and 12-LOX-/-platelets (dark grey bars) and plotted on logarithmic scale. **(C)** Individual PUFA concentrations tested after 24h of storage in WT (light grey bars) and 12-LOX-/-platelets (dark grey bars) and plotted on logarithmic scale. **(D)** Individual oxylipin concentrations tested after 24h of storage in WT (light grey bars) and 12-LOX-/-platelets (dark grey bars) and plotted on logarithmic scale. **(E)** Arachidonic acid supernatant concentration fresh, 24h, and 48h stored WT (light grey bars) and 12-LOX-/- (dark grey bars). (F) ω-6/ω-3 PUFA ratio (included as ω-6: AA, DGLA, GLA, LA, FA22_5w6, FA 22_4, included as ω-3: EPA, DHA, DPA, ALA) Results shown as mean ± SEM. * p<0.05, # p<0.01, $ p<0.001, @ p<0.0001 (no individual significances presented because of quantity).

### Platelet survival in vivo

In healthy humans, increased 12-HETE levels in stored platelets are associated with reduced platelet survival *in vivo*.^8^ To test how deletion of 12-LOX (the 12-HETE producing enzyme) affects platelet *in vivo* survival, we transfused fresh and stored WT and 12-LOX-/-platelets to WT mice. To distinguish between endogenous and transfused platelets, we biotinylated platelets with N-hydroxysuccimido (NHS) biotin *in vivo* based on a previously published protocol.^24^ We confirmed that the recovery and survival of fresh biotinylated platelets were indistinguishable from fresh UbiC-GFP platelets (Supplemental Figure 2). We obtained whole blood from mice after *in vivo* biotinylation to isolate platelet-rich plasma (PRP) (Figure 2A). PRP was transfused fresh after 24 or 48 hours of storage. We assessed platelet survival by phlebotomy after one, 24, 48, and 72 hours followed by staining with streptavidin and testing by flow cytometry. Unexpectedly, we found significantly reduced platelet survival with fresh 12-LOX-/-platelets compared to WT platelets at all time points (Figure 2B). However, stored 12-LOX-/- and WT platelets did not differ, either after 24 hours or 48 hours of storage at any time points tested (Figure 3C, D).

**Figure 2:**
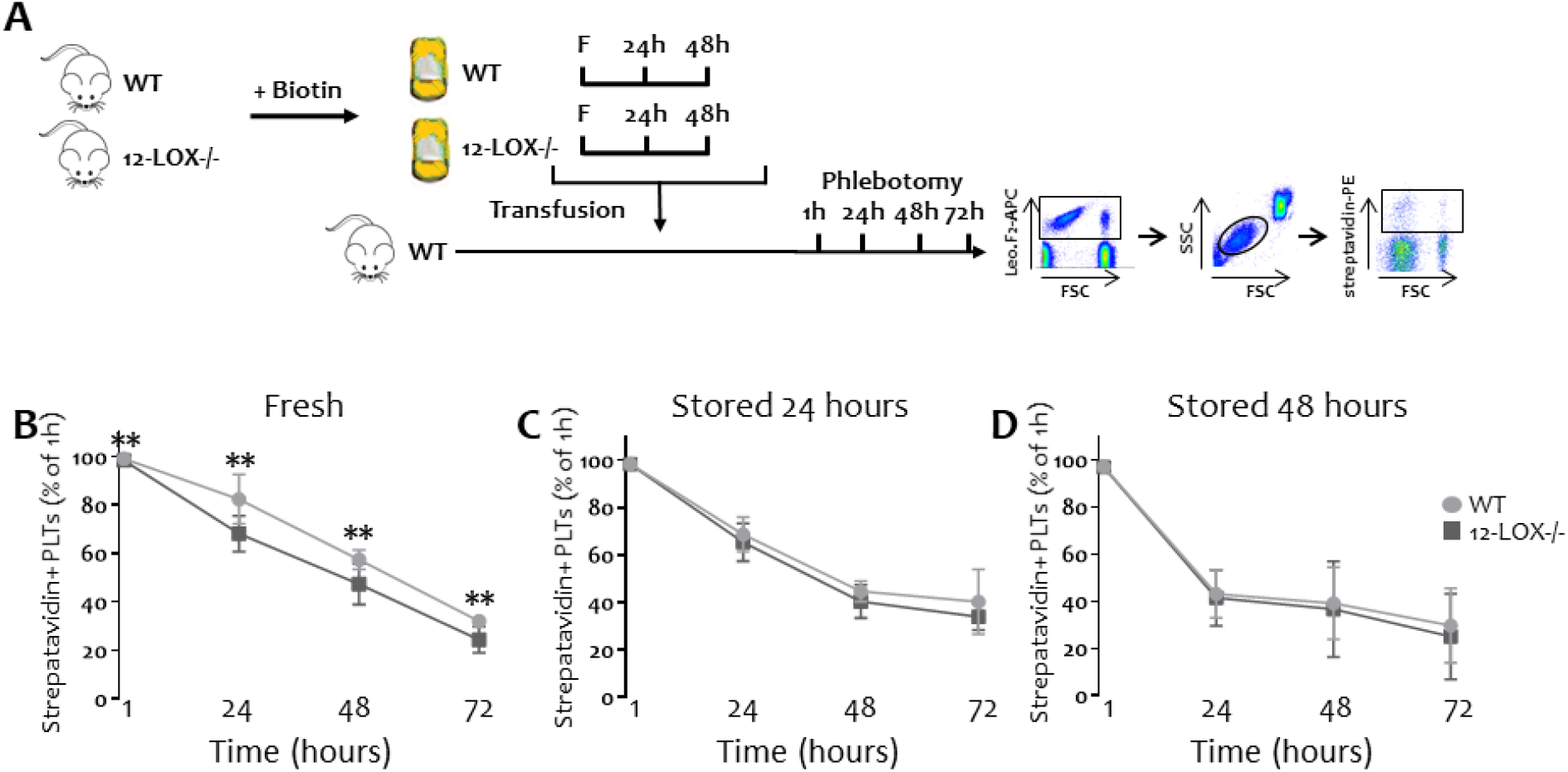
In vivo survival of 12-LOX-/- and WT platelets.

**(A)** Outline of experimental approach, Transfused WT platelets into WT mice (light grey circles), transfused 12-LOX-/-platelets into WT mice (dark grey squares). Survival of fresh **(B)**, 24 hour stored **(C)**, 48 hour-stored **(D)** WT and 12-LOX-/-platelets (streptavidin positive events by flow cytometry) over 72 hours, plotted as percentage of 1 hour count. Data shown as mean ± SEM, n=9, **p=0.00738 for 1 hour, **p= 0.00365 for 24 hours, **p=0.00512 for 48 hours, **p=0.00126 for 72 hours.

**Figure 3:**
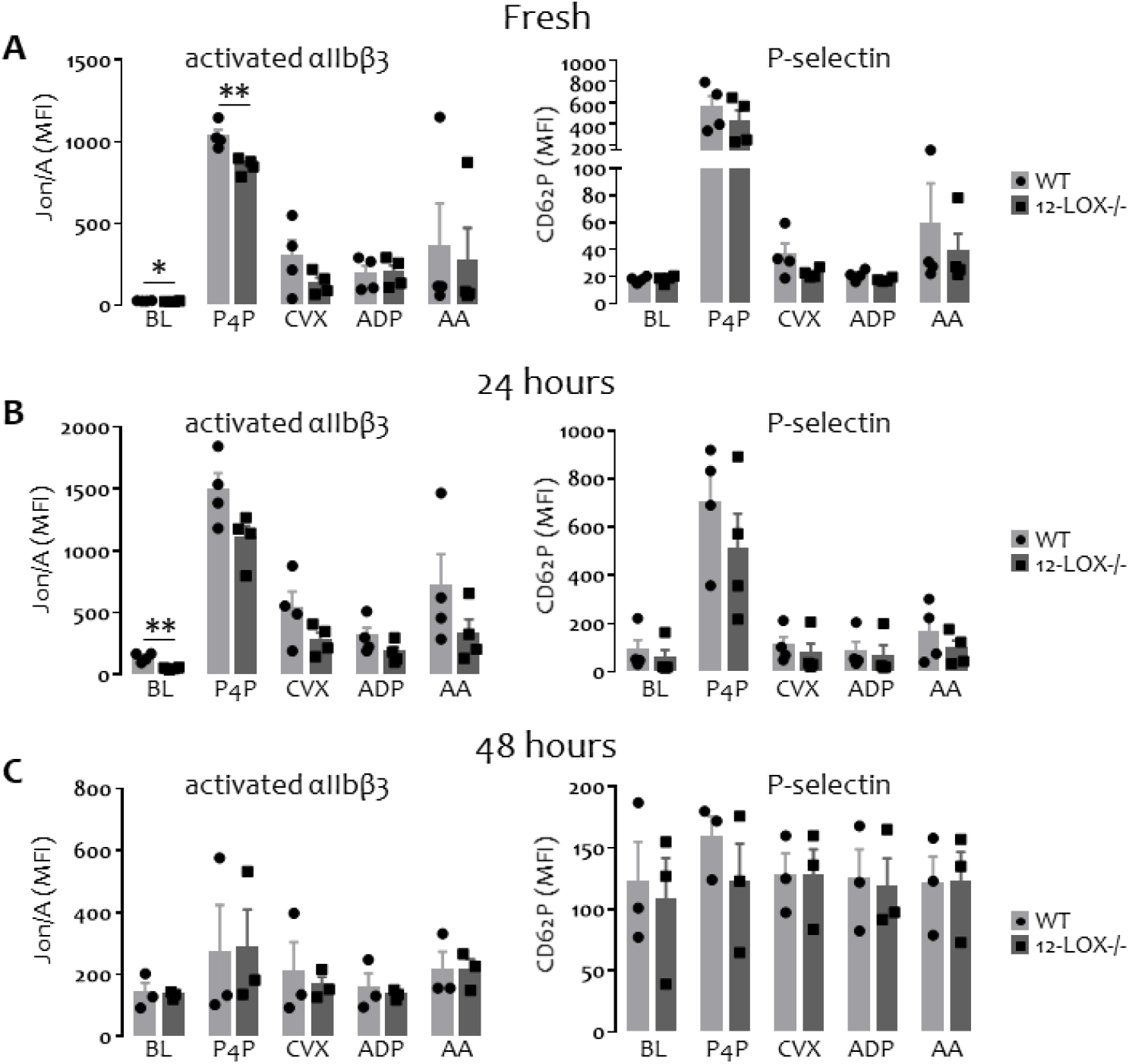
Pre-transfusion *in vitro* function of WT and 12-LOX-/-platelets.

PRP was stored at RT for 24 and 48 hours and aIIbb3 activation and α-degranulation was tested in response to different agonists (PAR-4 Peptide 0.5mM, Convulxin 100ng/mL, ADP 5uM, and Arachidonic Acid 0.5mM) for fresh, 24 hour-stored, and 48 hour-stored platelets. WT platelets (light grey bars with black circles), transfused 12-LOX-/-platelets (dark grey bars with black squares). Fresh **(A)**, 24 hour-stored **(B)**, 48 hour stored **(C)** WT and 12-LOX-/-platelet αIIbβ3 integrin function (left panel) and α-granule secretion (right panel) tested by flow cytometry. Samples were tested at baseline (BL), and after stimulation PAR-4 Peptide (P4P) 0.5mM, Convulxin (CVX) 100ng/mL, ADP 5uM, and Arachidonic Acid (AA) 0.5mM PAR-4. Data shown as mean ± SEM with individual data points, n=3-4, *p=0.0363, **p=0.0076 for fresh, P4p-treated platelets, **p=0.0026 for unstimulated (BL) 24 hour-stored platelets.

### Function of fresh and stored platelets before transfusion

Previous studies have described how 12-LOX deficiency affects platelet reactivity in fresh platelets.^5,27,28^ We corroborated these studies and found significantly decreased αIIbβ3 integrin activation in 12-LOX-/-platelets at baseline (Figure 3A). We found a trend for reduced αIIbβ3 integrin activation in response to convulxin and arachidonic acid. Responses to PAR-4 peptide were significantly lower in 12-LOX-/-platelets (Figure 3A). We detected no difference in response to ADP.

Similarly, we found a trend for reduced α-degranulation in response to all agonists, except at baseline and after stimulation with ADP. Storage is known to decrease platelet function.^29^ After 24 hours, we found more variability but a similar trend for decreased integrin activation and α-granule secretion and significantly lower baseline integrin activation in 12-LOX-/-platelets (Figure 3B). After 48 hours, the platelet response was universally lower, and no trends or significant differences were observed (Figure 3C). Conventional platelet storage parameters, including glucose, pH, morphology score, platelet counts, and mean platelet volume (MPV) did not differ significantly between WT and 12-LOX-/-platelets (Supplemental Figure 3). Our lipidomics data suggest that the maximum PUFA and oxylipin difference already occurs after 24 hours. We observed a significant pH decrease after 48 hours for both WT and 12-LOX-/-platelets (Supplemental Figure 3). We therefore included only the 24 hour storage time point for the subsequent *in vivo* studies.

### Function of fresh and stored platelets after transfusion in thrombocytopenic mice

The majority of stored platelets are transfused to prevent bleeding in thrombocytopenic patients.^30^ To test the post-transfusion function of WT and 12-LOX-/-platelets, we utilized a previously described murine thrombocytopenia model (Figure 4A). As previously described, we induced thrombocytopenia in human IL4 receptor-transgenic mice with an anti-human IL4-receptor antibody.^31^ We confirmed successful platelet depletion within 12 hours (Figure 4A). Because non-transgenic mouse platelets do not express the receptor for human IL4, we were able to reconstitute depleted mice with either fresh or 24-hour stored WT and 12-LOX-/-platelets (Figure 4A). One hour after transfusion of fresh platelets, we found no longer significantly lower αIIbβ3 integrin activation at baseline or in response to PAR-4 peptide (Figure 4B). Twenty-four hours after transfusion, we observed a trend for more platelet activation with WT platelets compared to 12-LOX-/-after stimulation with PAR-4 peptide approaching significance (p=0.07), similar to what we observed before transfusion with fresh platelets (Figure 4B). Surprisingly, when we transfused 24-hour stored platelets, baseline αIIbβ3 integrin activation was significantly higher in 12-LOX-/-platelets, in contrast to what we observed before transfusion (Figure 4C versus Figure 2B). Similarly, the trend for a decreased response to convulxin before transfusion was reversed, and we observed significantly more integrin activation after transfusion (Figure 4C).

**Figure 4:**
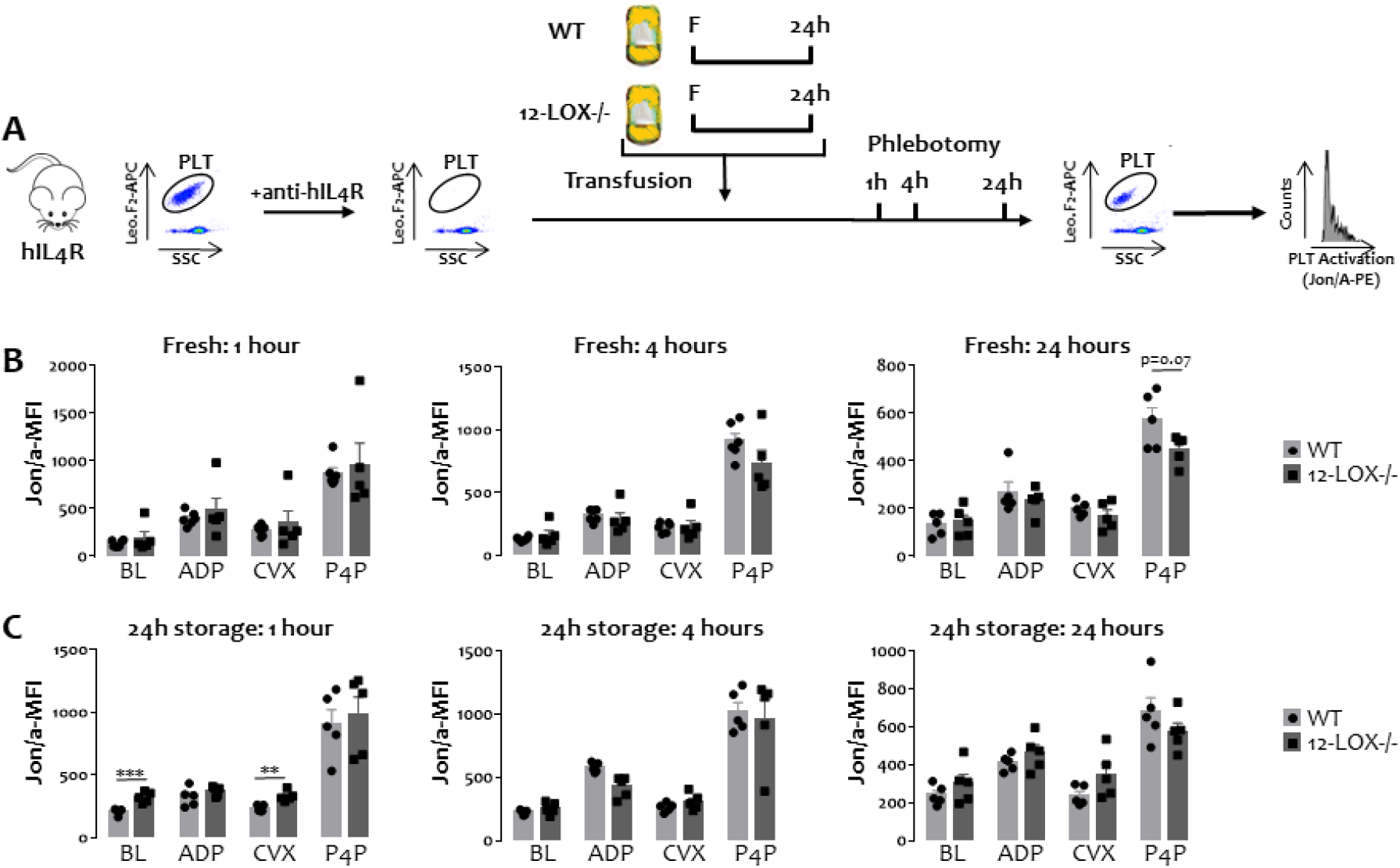
Post-transfusion function *in vitro* function of fresh and stored WT and 12-LOX-/-platelets.

PRP was stored at RT for 24 hours and 500uL platelets at 200 ×10^3^/μL were transfused into platelet-depleted hIL-4R-TG mice **(A)**. Mice were phlebotomized at 1, 4, and 24 hours post-transfusion and samples were analyzed by flow cytometry for platelet αIIbβ_3_ integrin function in response to different agonists (ADP 5uM, Convulxin 100ng/mL, and PAR-4 Peptide 0.5mM). Transfused WT platelets (light grey bars with black circles), transfused 12-LOX-/-platelets (dark grey bars with black squares). Fresh **(B)**, and 24 hour-stored **(C)** WT and 12-LOX-/-platelet αIIbβ_3_ integrin function tested one hour (left panel), four hours (middle panel), and 24 hours (right panel) post transfusion into thrombocytopenic mice, tested by flow cytometry. Data shown as mean ± SEM and individual data points, n=5-6, **p=0.00241, ***p=0.00093.

Over time, the αIIbβ3 activation in 12-LOX-/-platelets was attenuated, and after 4 and 24 hours, we did not find a significantly higher baseline αIIbβ3 increase anymore (Figure 4C).

### In vivo function of fresh and stored platelets after transfusion in thrombocytopenic mice

To test whether the increase in baseline integrin activation post-transfusion translates to increased *in vivo* function, we studied the functional response of WT and 12-LOX-/-platelets in ferric chloride (FeCl^3^)-injured carotid arteries, a macrovascular mouse thrombosis model as described before by us and others.^26^ In this model, thrombus formation in the carotid artery is detected by a reduction in blood flow rate. To focus on stored platelet quality and avoid interference with supernatant components, we washed stored platelets as described before transfusion. Similar to our post-transfusion platelet activation assay, we depleted hIL4R-TG mice with an anti-human hIL4R antibody and transfused 4×10^8^ 24-hour-stored WT or 12-LOX-/-platelets in 200μL. We phlebotomized recipient mice five minutes after transfusion to confirm an adequate count increase to allow for sufficient *in vivo* hemostasis of at least 100×10^3^/μL (Figure 5B).^32^ We observed no significant platelet count differences between hIL4R-TG mice transfused with WT or 12-LOX-/-platelets (Figure 5B). Confirming our prediction, the increased integrin activation in 12-LOX-/-platelets led to a significantly shorter time to occlusion in hIL4-TG mice transfused with 12-LOX-/-platelets (Figure 5C). The time to occlusion of transfused 12-LOX-/-mice was similar to WT mice with normal (unaltered) platelet counts (Figure 5C). In contrast, two out of five hIL4R-TG mice transfused with 24-hour stored WT platelets showed patent vessels until the experiment was terminated after 30 minutes. Overall, the time to occlusion was significantly longer with stored, transfused WT platelets than with stored, transfused 12-LOX-/-platelets or WT control mice (Figure 5C). Neither platelet depleted, untransfused hIL4R-TG mice nor hIL4-TG mice with normal platelet counts showed vessel occlusion in this model, highlighting the importance of platelet number and function in this model (hIL4R-TG mice lack functional GPIbα on the platelet surface) (Figure 5C).

**Figure 5:**
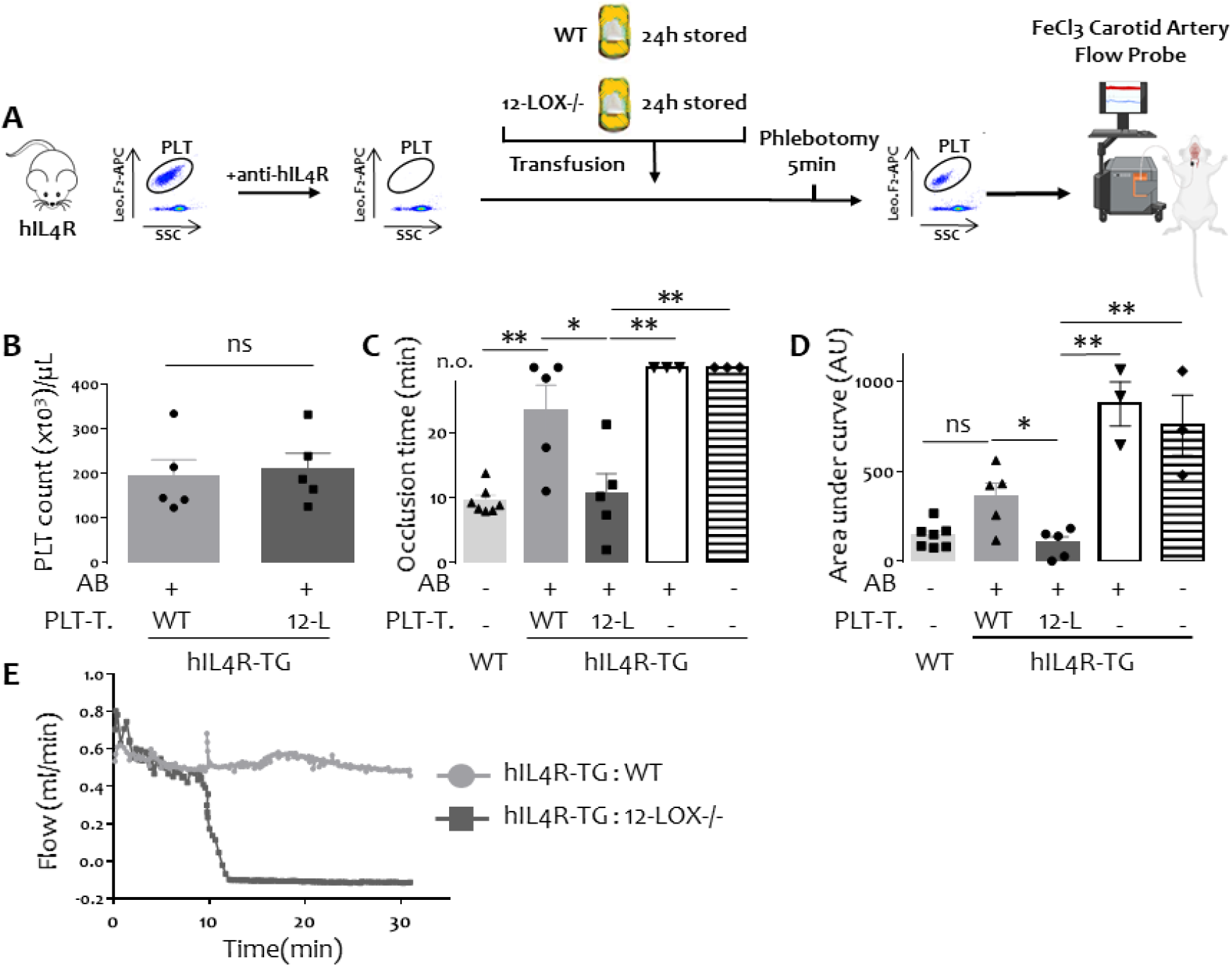
Post-transfusion *in vivo* function of WT and 12-LOX-/-platelets in thrombocytopenic mice.

Platelets were stored at RT for 24 hours, washed, and 4×10^8^ platelets were transfused into platelet-depleted hIL4R-TG mice. The carotid artery of an 8-10 week old male mouse was exposed to a 2M FeCl3 saturated strip of filter paper for 4 minutes, after which the artery was rinsed with PBS and the arterial flow rate was recorded for 30 minutes by flow probe. WT mice normal platelet counts (Off-white bars with black triangles), antibody depleted hIL4R-TG mice transfused with stored WT platelets (light grey bars with black circles), antibody depleted hIL4-TG mice transfused with stored 12-LOX-/-platelets (dark grey bars with black squares), antibody depleted hIL4-TG mice without transfusion (white bars with upside down triangles), hIL4R-TG mice with normal platelet counts (white-black-striped bars with black diamonds). *AB* indicates platelet depleting antibody injection (+/−), *PLT-T.* indicates stored platelet transfusion (platelet type / no transfusion [-]) **(A)** Overview of experimental outline **(B)** Post transfusion platelet count in platelet depleted hIL4R-TG mice **(C)** Occlusion time after FeCl3 injury in minutes defined as time from injury to cessation of blood flow (n.o. = no occlusion at termination of experiment), **(D)** Area under the curve (arbitrary units, AU) for 30min observation time, **(E)** Representative flow traces of hIL4R-TG mice transfused with WT (light grey circles) or hIL4R-TG platelets (dark grey squares). Data shown as mean ± SEM and individual data, n=3-7, ns = not significant, *p<0.05, **p<0.01.

In contrast, all mice transfused with 12-LOX-/-platelets showed vessel occlusion within 22 minutes (Figure 5C). Because a failure to occlude is only inadequately represented by the “time to occlusion” analysis, we measured the area under the curve (AUC) for the same experiments. Again, the AUC was significantly smaller in hIL4R-TG mice transfused with 12-LOX-/-platelets compared to those transfused with WT platelets proving significantly better platelet function *in vivo* (Figure 5D, E).

## Discussion

Our study has three major findings: 1) Storing platelets with a deletion of 12-LOX increases ω-6 and ω-3 PUFAs and decreases oxylipins compared to stored WT platelets. 2) Platelet survival is reduced in fresh but not in stored 12-LOX-/-platelets compared to WT platelets. 3) 12-LOX deletion decreases platelet activation in fresh and stored platelets *in vitro*, but upon transfusion into thrombocytopenic mice, 12-LOX-/-platelets show higher baseline integrin activation and improved *in vivo* function. Our findings are surprising because we predicted the opposite results, i.e., prolonged platelet survival and reduced *in vivo* function, based on previously published data.

Our targeted lipidomics results indicate significant changes between 12-LOX-/- and WT platelet samples. In WT platelets, 12-LOX catalyzes the oxygenation of ω-3 and ω-6 PUFAs after they are hydrolyzed from the membrane by PLA2. However, without 12-LOX activity, both ω-3 and ω-6 PUFAs accumulate, but our observed ω-6/ω-3 ratio decrease hints at a pronounced ω-3 PUFA increase.

Most studies on platelets with changes in PUFAs were performed in humans or mice on PUFA-rich diets. Ω-3 PUFAs offer protection from cardiovascular disease due to decreasing platelet aggregation and reduced thromboxane release.^33, 34^ Both EPA and DHA have been reported to inhibit platelet function, but DHA is much more potent.^35^ DHA-derived, 12-LOX-dependent oxylipins inhibit platelet reactivity in a glycoprotein VI-dependent manner via activation of protein kinase A.^19^ A diet rich in ω-3 PUFAs showed delayed platelet-dependent thrombin generation in both humans and mice *in vitro* and *in vivo* leading to a reduced thrombus size and delayed vessel occlusion, without affecting the bleeding time.^36^ The results of our *in vitro* (pre-transfusion) data match the studies outlined above. However, the relevance of different PUFA compositions for stored platelets and post-transfusion rejuvenation has never been studied. Based on our data, a diet rich in fish oil could yield a better platelet product for transfusion. One caveat is that the platelet count of healthy humans on fish oil supplementation is mildly but significantly decreased.^37-39^ This could suggest that increased ω-3 PUFAs lead to increased clearance, e.g., by increased sequestration due to increased membrane fluidity. Therefore, the functional increase could come at the expense of mildly reduced platelet survival. In our study, we observed decreased platelet survival mainly in fresh 12-LOX-/-platelets that showed an isolated increase in the ω-3 PUFAs EPA and DPA and a significantly lower ω-6/ω-3 PUFA ratio than WT platelets. We no longer observed this phenomenon in stored platelets, possibly due to the general detrimental nature of the storage lesion or the significant increase in all ω-3 *and* ω-6 PUFAs. Whether ω-6 PUFAs have a prolonging effect on platelet circulation time has never been tested but is another possible explanation for our findings. How incomplete inhibition of 12-LOX, e.g., by pharmacological means, affects ω-3 and ω-6 PUFAs and oxylipins during storage and after transfusion will have to be studied in the future.

The most striking finding of our study is the improvement of stored platelet function after transfusion *in vivo*. We are unaware of other studies showing that the storage lesion can be bypassed with the same level of solid post-transfusion, in vivo evidence. After transfusion, the increase ω-6 PUFAs (especially arachidonic acid) led to a significant increase in platelet function, and vessel occlusion occurred at a similar rate as in unaltered, healthy WT mice.

In contrast to PUFAs, oxylipins were depleted in 12-LOX-/-platelets. It could be assumed that PUFAs would be routed through other enzymatic pathways if 12-LOX is inactive, such as other LOX enzymes or COX, but that does not appear to be the case. The most significant changes between 12-LOX-/- and WT platelets occurred in 12-HETE, 12-HEPE, and 14-HDoHE, which are the main oxylipins produced by 12-LOX from AA, EPA, and DHA, respectively. Interestingly, we also found that 15-LOX and COX metabolites were significantly decreased in 12-LOX-/-platelets. This is likely because unlike in humans, where 15-LOX1 and 15-LOX2 are separated from 12-LOX, in the mouse 12-LOX (ALOX12) is a 12/15-LOX that produces both 12 and 15 oxidation products.^1^ However, there is a 6-10:1 ratio in favor of 12 oxygenation products relative to 15 oxygenation products.^40^

Previous studies have altered the arachidonic acid metabolism by inhibiting cyclooxygenase-1 (COX-1) with acetylsalicylic acid (ASA, aspirin) in stored platelets. While platelet aggregation was inhibited before storage compared to the same group without ASA, no differences were observed post storage, suggesting that there is no detrimental effect after storage.^41^ Similar to our study, conventional storage parameters, such as pH, LDH, lactate, and CD62P did not differ between the two groups. Whether there are differences after transfusion (i.e., differences in post-transfusion rejuvenation) has not been studied. Adding ASA to stored platelets also did not affect GPIbα shedding or CD40L level.^42^ In contrast to 12-LOX, inhibiting COX-1 with ASA blocks thromboxane generation before and after transfusion and is thereby more likely to *inhibit* instead of promoting platelet function, even post-transfusion.

Therefore, a possible explanation for our findings is that once stored 12-LOX-/-platelets are transfused and thereby rejuvenate *in vivo*, the accumulated arachidonic acid is metabolized predominantly through COX-1 to thromboxane. This hypothesized mechanism explains why we observed reduced integrin activation and function before transfusion and the opposite after transfusion. Following this hypothesis, one would predict that by inhibiting COX-1, platelets would metabolize their accumulated arachidonic acid predominantly through 12-LOX and yield increased 12-HETE. This is likely unfavorable given 12-HETE’s role in TRALI, inflammation, and reduced platelet survival in healthy humans. However, this will have to be investigated in future studies. Recent clinical trials suggest a critical unmet need for improved function in stored platelets. In neonates, a higher transfusion threshold worsened outcomes and increased bleeding events.^9^ Similarly, patients with intracranial hemorrhage had worse outcomes when they received platelets compared to supportive therapy without platelet transfusions.^10^

In summary, our data suggest that inhibition of 12-LOX in stored platelets could be a promising therapeutic approach to improve stored platelet function for therapeutic and prophylactic transfusions.

## Acknowledgements

The authors would like to thank Dr. Jerry Ware from the University of Arkansas for providing hIL4R-TG mice, and Renetta Stevens, Tena Petersen for administrative support. M.S. received funding from the American Heart Association (Career Development Award), NIH (1R01HL153072-01), and DoD (W81XWH-12-1-0441, EDMS 5570).

## Authorship contributions

H.J. and D.B. performed most experiments, analyzed data and wrote the manuscript, S.L.B. designed and performed experiments and analyzed data, M.C.S. performed experiments and analyzed data, M.H. provided mice, analyzed data, and wrote the manuscript, X.F. designed experiments, analyzed data, and wrote the manuscript, M.S. outlined the study, designed and performed experiments, analyzed data, and wrote the manuscript

---

**Supplemental Figure 1:**
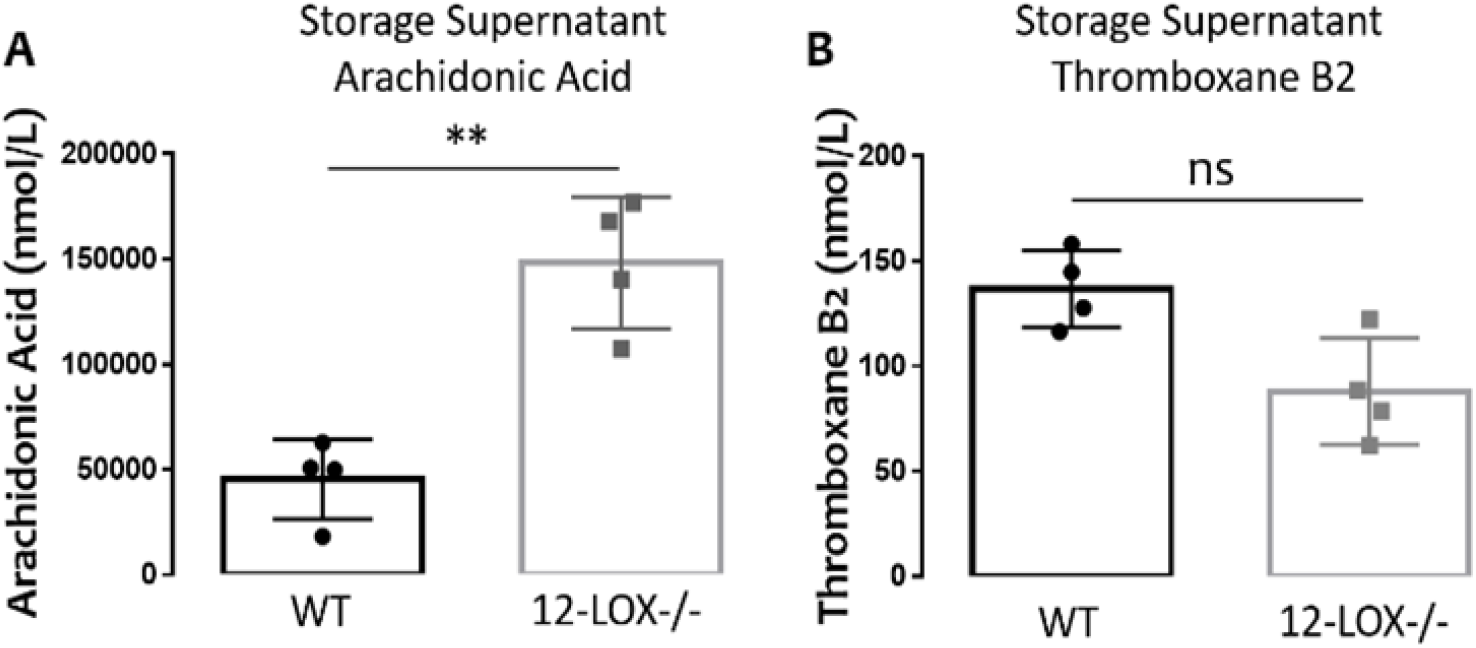
Stored supernatant from 24 hour-stored WT (black outline with circles) and 12-LOX-/- (grey outlines with squares) platelets was analyzed for arachidonic acid (A) and thromboxane B2 (B) by targeted LC-MS/MS. Results shown as mean ± SEM, n=4, **p=0.0013

**Supplemental Figure 2:**
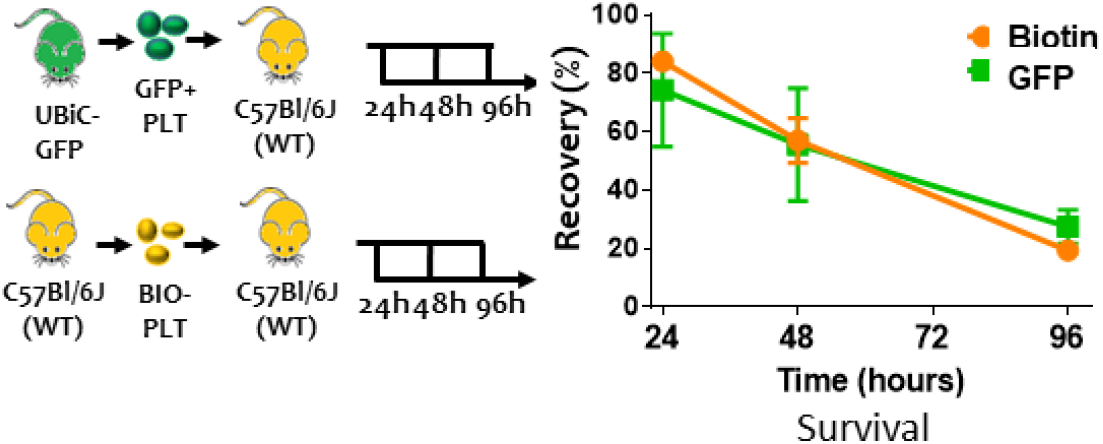
Comparison of *in vivo* survival of UbiC-GFP platelets and biotin-labeled WT mice. UbiC-GFP platelets (green squares) or biotin positive platelets (orange circles) were isolated and transfused to WT mice and samples were obtained after 24, 48, 72, and 96 hours to trace GFP positive or streptavidin-PE positive events by flow cytometry.

**Supplemental Figure 3:**
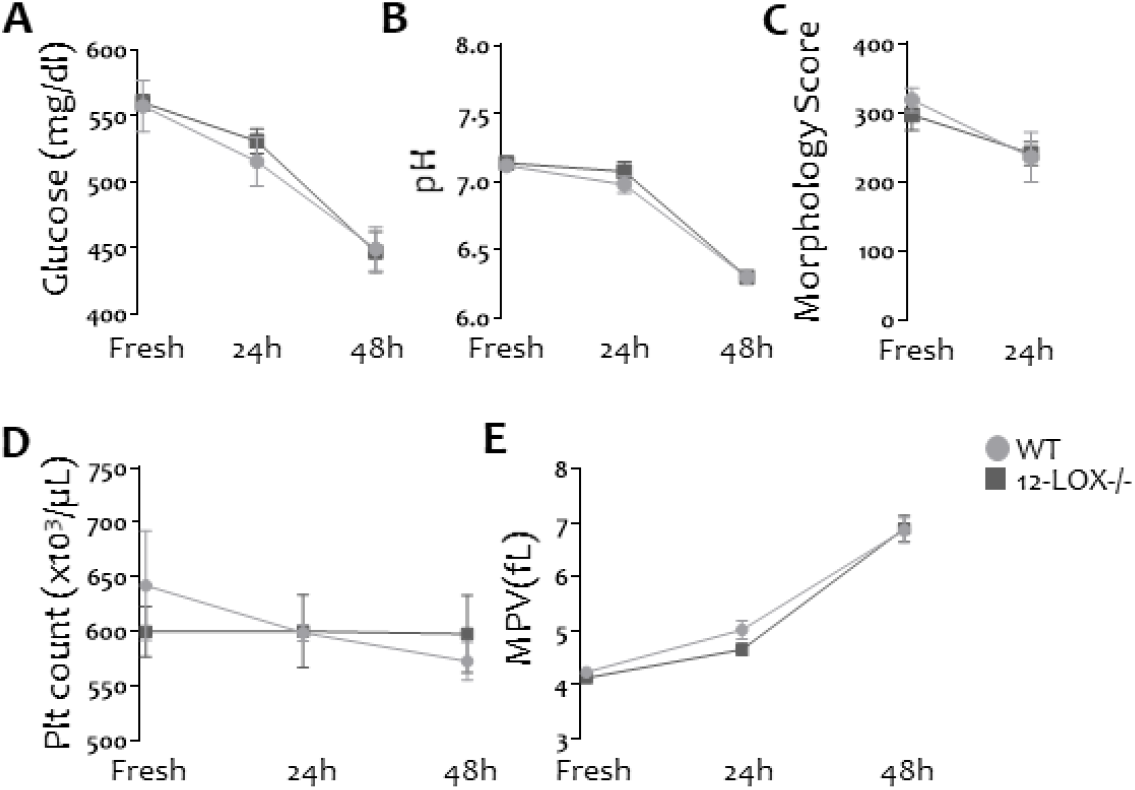
Comparison of conventional *in vitro* storage parameters of WT platelets and 12-LOX-/-platelets.

WT platelets (light grey circles) or 12-LOX-/-platelets (dark grey squares) were isolated and tested fresh, after 24, and 48 hours of storage. **(A)** Glucose, and **(B)** pH was measured by Blood Gas Analzyer, **(C)** Kunicki’s Morphology Scores was assessed by light microscopy, **(D)** Platelet count and **(E)** MPV were tested by ABX Hemanalyzer. Data are shown as mean ± SEM. N=4-27

## Notes

C.O.I.: M.S. received research funding from Terumo BCT and Cerus Corp. All other authors have no conflict of interest to declare

### Competing Interest Statement

The authors have declared no competing interest.

